# Cysteine tRNA acts as a stop codon readthrough-inducing rti-tRNA in the human HEK293T cell line

**DOI:** 10.1101/2023.04.17.537261

**Authors:** Leoš Shivaya Valášek, Michaela Kučerová, Jakub Zeman, Petra Beznosková

## Abstract

Under certain circumstances, any of the three termination codons can be read through by its near-cognate tRNA; i.e. a tRNA whose two out of three anticodon nucleotides base-pair with those of the stop codon. Unless programmed to synthetize C-terminally extended protein variants with expanded physiological roles, readthrough represents an undesirable translational error. On the other side of a coin, a significant number of human genetic diseases is associated with the introduction of nonsense mutations (premature termination codons - PTCs) into coding sequences, where stopping is not desirable. Here, the tRNA’s ability to induce readthrough opens up the intriguing possibility of mitigating the deleterious effects of PTCs on human health. In yeast, the UGA and UAR stop codons were described to be read through by four readthrough-inducing rti-tRNAs – tRNA^Trp^ and tRNA^Cys^, and tRNA^Tyr^ and tRNA^Gln^, respectively. The readthrough-inducing potential of tRNA^Trp^ and tRNA^Tyr^ was also observed in human cell lines. Here, we investigated the readthrough-inducing potential of human tRNA^Cys^ in the HEK293T cell line. The tRNA^Cys^ family consists of two isoacceptors, one with ACA and the other with GCA anticodons. We selected nine representative tRNA^Cys^ isodecoders (differing in primary sequence and expression level) and tested them using dual luciferase reporter assays. We found that at least two tRNA^Cys^ can significantly elevate UGA readthrough when overexpressed. This indicates a mechanistically conserved nature of rti-tRNAs between yeast and human, supporting the idea that they could be utilized in the PTC-associated RNA therapies.

## INTRODUCTION

tRNA molecules are non-coding RNA molecules, usually ∼73-93 nucleotides long, carrying a specific amino acid attached to their 3’ OH end. In this so-called charged aminoacyl-tRNA (aa-tRNA) state, they form an elongation ternary complex with eEF1A and GTP, whose role is to deliver aa-tRNAs to the ribosomal A-site during translation elongation. Upon sense codon recognition, GTP on eEF1A is hydrolyzed triggering the release of eEF1A-GDP from the ribosome and subsequent accommodation of the aa-tRNA in the decoding site to proceed to the peptide bond formation. Once one of the three stop codons occurs in the A-site, a release ternary complex consisting of eRF1/eRF3-GTP enters the decoding site to recognize the stop (Hellen 2018). Analogously, this triggers GTP on eRF3 to be hydrolyzed and released, allowing the catalytic GGQ motif of eRF1 to reach to the peptidyl-transferase center next to the 3’ CCA end of the peptidyl-tRNA sitting in the ribosomal P-site to catalyze the peptide release.

When the elongation complex with aa-tRNA, which is near cognate to a stop codon occurring in the A-site, enters it prior to the release eRF1/eRF3-GTP complex, translational stop codon readthrough (SC-RT) can occur instead of termination. The near-cognate tRNAs (nc-tRNAs) are characterized by having two of the three anticodon nucleotides complementary to a given stop codon. Simply speaking, when SC-RT occurs, the stop codon is redefined as sense; i.e. an additional amino acid is incorporated to the polypeptide chain and translation continues until the next stop codon, which leads to production of the C-terminally extended proteins (Dabrowski et al. 2015). Expectedly, a natural/spontaneous frequency of translational (basal) SC-RT is quite low, < 0.1 % (Keeling et al. 2012). It can be influenced by the identity of the stop codon, nucleotide context, *cis*-acting mRNA elements and trans-acting factors (Skuzeski et al. 1991; Bonetti et al. 1995; McCaughan et al. 1995; Cassan and Rousset 2001; Namy et al. 2001; Harrell et al. 2002; Firth et al. 2011; Floquet et al. 2012; Beznoskova et al. 2015; Beznoskova et al. 2016; Gunisova et al. 2016; Beznoskova et al. 2019). Naturally occurring SC-RT can also be programmed/desired, producing C-terminally extended proteins with new biological roles (Yamaguchi et al. 2012; Eswarappa et al. 2014; Schueren et al. 2014; Loughran et al. 2018). Intriguingly, the extreme case of SC-RT is the reassignment of all three stop-codons as sense codons in some ciliates and trypanosomatids (Heaphy et al. 2016; Swart et al. 2016; Zahonova et al. 2016; Kachale et al. 2023). It is highly likely that programmed SC-RT is affected by similar factors as the basal one, however, what factors create the major difference only remains to be seen (Martins-Dias and Romao 2021).

Several naturally occurring tRNAs can work as nc-tRNAs. The mismatch occurs most frequently at position 1 or 3 of a stop codon. The list of all possible nc-tRNAs with these mismatches is given in Table 1. In yeast, it was shown that UAA and UAG stop codons can be reassigned with Gln, Lys or Tyr, while the UGA stop codon with Trp, Arg or Cys (Table 1) (Blanchet et al. 2014; Roy et al. 2015; Blanchet et al. 2018). In particular, Tyr and Gln were inserted at much higher frequency at UAA than Lys. Gln was also the most frequent amino acid inserted at UAG, with Tyr and Lys showing much lower frequencies. Interestingly, no Glu was observed to be inserted at UAA or UAG. As for the UGA stop codon, there exits two types of nc-tRNAs with a mismatch at the 3^rd^ position (tRNA^Trp^[CCA] and tRNA^Cys^[GCA or ACA]), and two with a mismatch at the 1^st^ position (tRNA^Arg^[UCU] and tRNA^Gly^[UCC]) (Table 1). Nonetheless, it was shown that UGA is decoded by nc-tRNAs with the wobble base-pair at the 3^rd^ position in most cases (Beznoskova et al. 2015; Roy et al. 2015; Beznoskova et al. 2019), with Trp being inserted at a higher frequency than Cys. Using well-established dual luciferase reporters, we systematically examined all nc-tRNAs for their ability to promote SC-RT in yeast and defined altogether four so-called readthrough-inducing tRNAs (rti-tRNAs) with high readthrough-inducing potential; namely tRNA^Trp^, tRNA^Tyr^, tRNA^Cys^, and tRNA^Gln^ (Beznoskova et al. 2016; Beznoskova et al. 2019). The former two were also tested in human cells with the same outcome (Beznoskova et al. 2021).

**Table 1.**
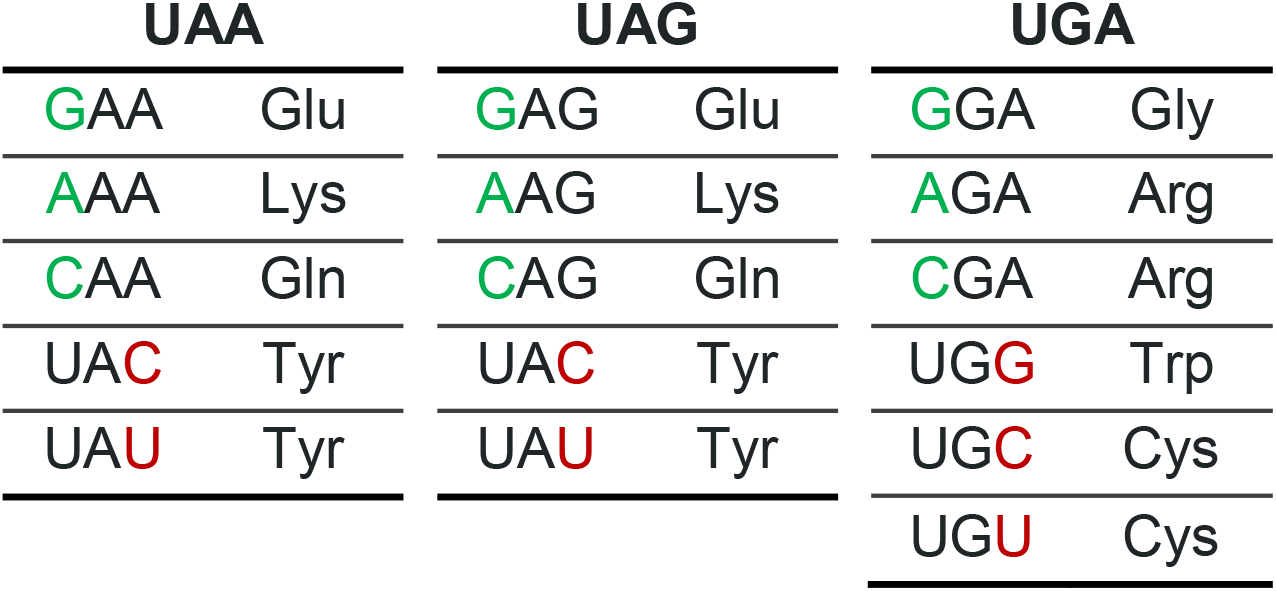
The list of codons with mismatch at position 1 or 3 of the UAA/UAG/UGA stop codons (eukaryotic genome). The inserted amino acids are indicated.

| UAA |  | UAG |  | UGA |  |
| --- | --- | --- | --- | --- | --- |
| GAA | Glu | GAG | Glu | GGA | Gly |
| AAA | Lys | AAG | Lys | AGA | Arg |
| CAA | Gln | CAG | Gln | CGA | Arg |
| UAC | Tyr | UAC | Tyr | UGG | Trp |
| UAU | Tyr | UAU | Tyr | UGC | Cys |
|  |  |  |  | UGU | Cys |

While spontaneous SC-RT at natural stop codons is not necessarily a desirable event, if nonsense mutations – so-called premature termination codons (PTCs) – occur in the coding regions of life-sustaining genes, induced SC-RT at these PTCs appears to be the best solution to deal with this problem (Keeling et al. 2014; Martins-Dias and Romao 2021). Naturally, PTCs abort translation elongation producing truncated protein variants that can have either gain-of-function or dominant-negative properties. In addition, mRNAs carrying PTCs become substrates of nonsense-mediated mRNA decay, which rapidly degrades them (Kurosaki and Maquat 2016). It was estimated that more than 15% of all human genetic diseases can be attributed to the presence of PTCs (Linde and Kerem 2008; Kurosaki and Maquat 2016). PTCs were shown to account for example for cystic fibrosis, spinal muscular atrophy (SMA), Duchenne muscular dystrophy or haemophilia. The search for an effective agent to specifically induce SC-RT in PTCs to prevent NMD and at least partially restore full-length expression of the affected protein has long been a major focus of this type of research. So-called readthrough inducing drugs (RTIDs) have become the most promising direction, but they still face drawbacks such as high toxicity and unpredictable efficacy in individual patients (Lee and Dougherty 2012; Keeling et al. 2014). Therefore, a second, much less explored direction focusing on the use of nc-tRNAs has recently gained increasing attention (see for example (Beznoskova et al. 2021)).

Here, we extended our analysis of rti-tRNAs in humans. Having recently demonstrated that tRNA^Trp^ boosts SC-RT at UGA and tRNA^Tyr^ at UAG and UAA in human cell lines (Beznoskova et al. 2021), here we focused on tRNA^Cys^, whose yeast counterpart belongs to a group of four well-characterized rti-RNAs (Beznoskova et al. 2016; Beznoskova et al. 2019). Using our dual luciferase systems coupled with the U6 promotor-based tRNA overexpression system, the nine tRNA^Cys^ with the highest mature tRNA score as predicted by tRNAscan-SE and/or the highest expression were subcloned into our expression vectors and tested for their ability to promote SC-RT in human HEK293T cells. Two of them, those with the best correlation of these two parameters, scored positively; therefore we report here the extension of the human rti-tRNA family with tRNA^Cys^.

## RESULTS

### Rational behind the selection of tRNA^Cys^ isodecoders from bioinformatically predicted human tRNA^Cys^-encoding genes

The tRNA^Cys^ family consists of two isoacceptors, one with ACA and the other with GCA anticodons. There is 1 isodecoder falling under ACA isoacceptors and 23 isodecoders falling under GCA isoacceptors. Considering that the mature sequences of all these isodecoders differ only slightly (see below), yet these nucleotide changes may play an important role in the overall tRNA expression levels and/or the SC-RT-promoting potential, we aimed to make a preselection of tRNA^Cys^ isodecoders to cover all their existing forms as closely as possible. The sequences of tRNA^Cys^ isodecoders were taken from GtRNAdb 2.0 (Chan and Lowe 2016), which uses the tRNAscan-SE 2.0 search tool to detect and functionally predict tRNA genes and annotate them in genomes (Lowe and Chan 2016). Since all these tRNAs are only bioinformatically predicted, it is possible that some of the isoforms may not be highly or even at all expressed *in vivo*.

When analyzing the tRNA^Cys^ isodecoders, the mature tRNAscan-SE score was considered the first. When higher than 50, it implies that the given tRNA most likely forms a functional secondary structure (Pan 2018). In support, a recent tRNA-Seq study revealed that tRNAs from HEK293T cells with tRNAscan-SE score >55 contributed the most to the total tRNA pool (Torres et al. 2019). All the 23 tRNA^Cys^[GCA] isodecoders have the mature tRNA score > 50 (Supplementary Table 1). The ACA isoacceptor (and the only isodecoder) has the score 46.2, its gene carries an intron inside the sequence, it is classified as pseudogene and, therefore, it was excluded from our analysis.

Next parameter that we considered was the “rank of isodecoder” section in GtRNAdb, which shows how high a particular tRNA^Cys^[GCA] scores relative to all other isodecoders in the same isodecoder family (Chan and Lowe 2016). Actually, this rank can be told directly from a gene name, for example with tRNA^Cys^[GCA]-1-1 ranking at the very top based on this parameter.

To assess the expression of tRNA^Cys^ isodecoders, three publicly available expression datasets were analyzed. The analysis of Solexa Illumina HiSeq 2000 small RNA-Seq gene counts in HEK293T cells (accession no. GSE114904, (Torres et al. 2019); Supplementary Table 2); the analysis of the data from Polymerase III chromatin immunoprecipitation followed by ChIP-Seq from liver cancer cell lines HepG2, Huh7 and healthy human liver (accession no. E-MTAB-958, (Rudolph et al. 2016), Supplementary Table 3); and the analysis of YAMAT-seq (Y-shaped Adapter-ligated MAture tRNA sequencing) in breast cancer cell lines MCF-7, SK-BR-3 and BT-20 (accession no. SRP096584; (Shigematsu et al. 2017), Supplementary Tables 4 - 5). The tRNA gene counts expressed as RPM (reads per milion) were divided into five groups according to their expression relative to the whole dataset, with 100% representing the isodecoder with the highest expression values. These were then classified into five groups marked with a plus sign; “+” meaning 0-20 %; “++”_20-40 %; “+++”_40-60 %; “++++”_60-80 %; “+++++”_80-100 %. The results were then cross-compared for all the three datasets and the highest scoring isodecoders were selected to be further tested experimentally. Table 2 summarizes the isodecoders selected for further study according to this analysis. A full list of all analysed isodecoders and their expression scores can be found in Supplemetary Table 6. The sequences of all tRNA^Cys^ isodecoders are aligned in Supplementary Figure S1. Note that the expression of isodecoder tRNA genes relative to their whole isodecoder set (based on the percentage of sequencing reads) was bioinformatically evaluated by applying a numerical analysis, isodecoder-specific tRNA gene contribution profile (iso-tRNA-CP), to several tRNA-Seq datasets from HEK293T cells. Overall, this showed, consistently with our analysis, that tRNA^Cys^[GCA-4-1] (Cys01) and tRNA^Cys^[GCA]-2 (Cys02) contribute the most to the tRNA^Cys^ isodecoder dataset (Table 2). Iso-tRNA-CP also showed that tRNA expression is cell-type and tissue specific, comparing tRNA-Seq datasets from the HEK293T cells and human brain (Torres et al. 2019).

**Table 2.** The tRNA-Cys isodecoders selected by the expression analysis that were selected for further analysis. The table shows the proportion of individual tRNA-Cys isodecoders expression relative to the whole isodecoder set of tRNA-Cys-GCA. ND – Not detected. The range of expression is indicated by a plus sign (“+”), “+++++” meaning the highest expression. The Cys02 sequence can be found in the human genome for four identical gene copies (tRNA-Cys-GCA-2-1, tRNA-Cys-GCA-2-2, tRNA-Cys-GCA-2-3, tRNA-Cys-GCA-2-4), the same applies for the Cys03 sequence (tRNA-Cys-GCA-9-1, tRNA-Cys-GCA-9-2, tRNA-Cys-GCA-9-3, tRNA-Cys-GCA-9-4).

| tRNA | Gene name | Tores, 2019 | Kutter, 2016 |  |  | Shigematsu, 2017 |  |  |
| --- | --- | --- | --- | --- | --- | --- | --- | --- |
|  |  | HEK293T | HepG2 | Huh7 | Liver-Adult | MCF-7 | SKBR-3 | BT-20 |
| Cys01 | tRNA-Cys-GCA-4-1 | + | ++++ | +++++ | ++++ | +++ | + | ++ |
| Cys02 | tRNA-Cys-GCA-2-1 | +++ | ND | ND | ND | +++++ | +++++ | ++++ |
|  | tRNA-Cys-GCA-2-2 | +++++ | ++ | +++ | ++ |  |  |  |
|  | tRNA-Cys-GCA-2-3 | + | +++++ | +++++ | +++++ |  |  |  |
|  | tRNA-Cys-GCA-2-4 | ++ | ND | ND | ND |  |  |  |
| Cys03 | tRNA-Cys-GCA-9-1 | + | ND | ND | ND | + | + | + |
|  | tRNA-Cys-GCA-9-2 | ++ | + | + | + |  |  |  |
|  | tRNA-Cys-GCA-9-3 | ++ | ND | ND | ND |  |  |  |
|  | tRNA-Cys-GCA-9-4 | ++ | + | + | + |  |  |  |
| Cys04 | tRNA-Cys-GCA-8-1 | + | + | + | + | + | + | + |
| Cys05 | tRNA-Cys-GCA-5-1 | ++ | + | + | + | ++++ | + | ++ |
| Cys06 | tRNA-Cys-GCA-14-1 | + | +++ | ++++ | +++ | + | + | + |
| Cys07 | tRNA-Cys-GCA-22-1 | + | ND | ND | ND | ND | ND | ND |
| Cys08 | tRNA-Cys-GCA-17-1 | +++++ | +++ | +++++ | ++ | ND | ND | ND |
| Cys09 | tRNA-Cys-ACA-1-1 | + | ND | ND | ND | ND | ND | ND |

### tRNA^Cys^ isodecoders with the highest tRNA mature score promote SC-RT on UGA when overexpressed

Each tRNA^Cys^ isodecoder was subcloned into the U6 promotor-driven tRNA overexpression cassette that is individually coupled with two basic dual luciferase p2luci and pSGDluc systems published before (Grentzmann et al. 1998; Loughran et al. 2017), where either the CAA sense or one of the stop codons separates two in-frame luciferases. These “coupled” constructs were created and thoroughly tested before (Beznoskova et al. 2021) (see also Methods), because this arrangement ensures that the measured readthrough values originated only from cells that did contain an ectopically expressed tRNA, reflecting their true capacity to boost stop codon recoding. For the sake of clarity, we first investigated if overexpression of any of the selected isodecoders coupled with the sense control p2luci plasmid affects the Firefly to *Renilla* ratio in the HEK293T cells. As shown in Supplementary Figure S2, we recorded comparable values of F/R ratio for all tested tRNAs^Cys^, as well as for the reporter with no tRNA (for numbers see Supplemental Excel File 1). These control experiments clearly demonstrated that neither of the tRNA^Cys^ isodecoders has any influence over the expression of both reporter genes when expressed under the U6 promoter. Similar results were previously observed for both tRNA^Tyr^ and tRNA^Trp^ (Beznoskova et al. 2021). These results entitled us to use the p2luci-CAG-no_tRNA vector as our only sense codon control for the subsequent experiments.

Next, we measured the SC-RT potential of all selected isodecoders setting the value obtained with the control p2luci-CAG-no_tRNA as 100 % efficiency. Based on our results, three groups of tRNAs^Cys^ isodecoders emerged: i) overexpression of Cys01 and Cys02 significantly increased SC-RT of UGA (by ∼ 6.0 % [P < 0.01] and ∼ 4.9 % [P < 0.05], respectively, compared to ∼ 3.1 % for the reporter with no tRNA) (Fig. 1A; Supplementary Table 1; for numbers see Supplemental Excel File 1); ii) overexpression of Cys03, Cys04 and Cys05 increased UGA SC-RT but only modestly (by ∼ 4.5 % [P < 0.1], ∼ 4.2 % [no significance], and ∼ 4.0 % [P < 0.1], respectively) (Fig. 1B; Supplementary Table 1; for numbers see Supplemental Excel File 1); and iii) overexpression of mature tRNA sequences of Cys06, Cys07, Cys08 and Cys09 did not change the UGA SC-RT levels at all (Fig. 1C; Supplementary Table 1; for numbers see Supplemental Excel File 1). We conclude that the degree of SC-RT efficiency perfectly correlates with the mature tRNA scores (as predicted by tRNAscan-SE); tRNAs^Cys^ with the highest scores displayed the most efficient SC-RT on UGA, the isodecoders displaying only a mild increase in SC-RT efficiency had lower scores. Notably, the same correlation was previously observed for tRNA^Tyr^ and tRNA^Trp^ (Beznoskova et al. 2021).

**Figure 1.**
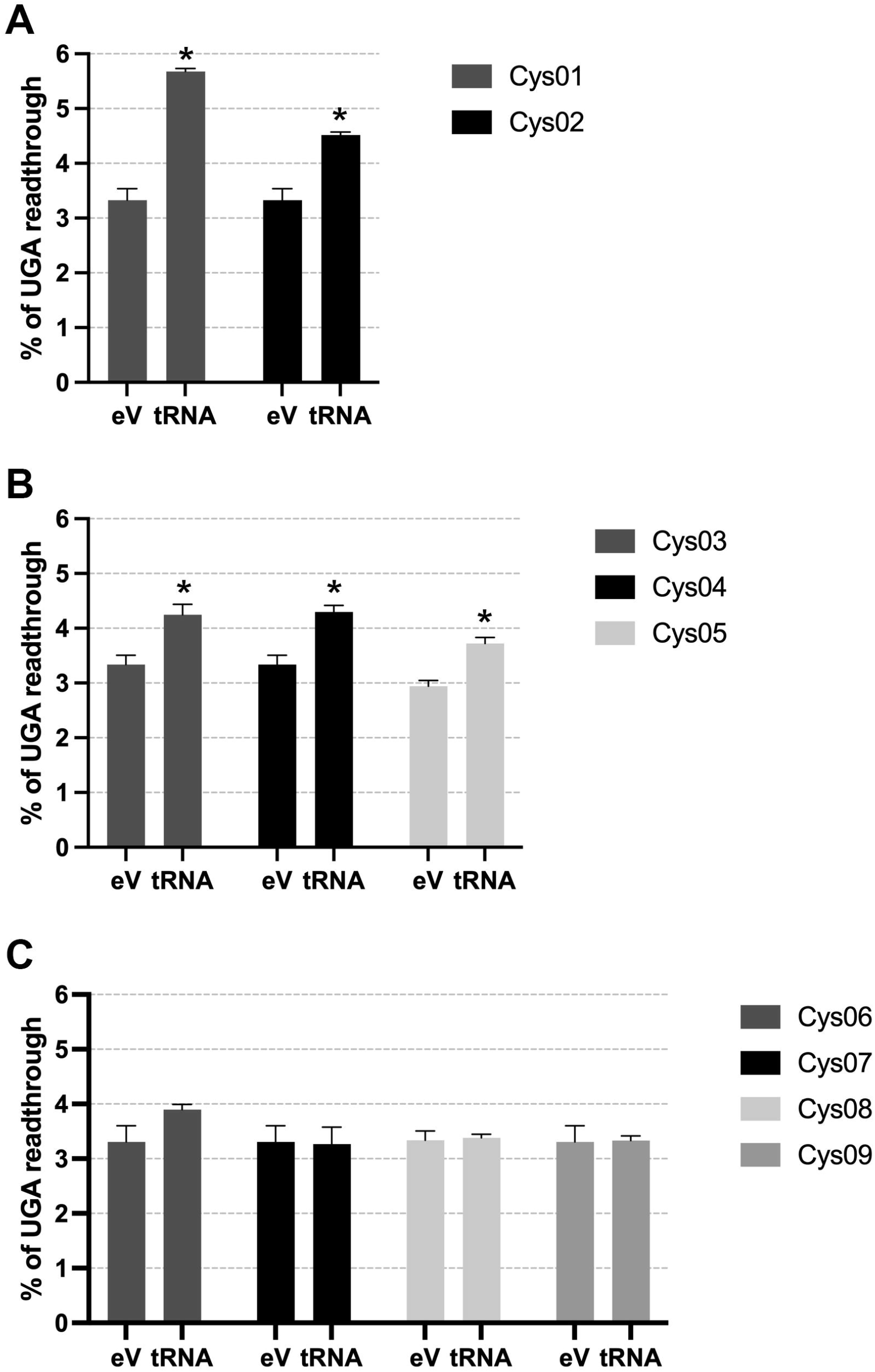
tRNA^Cys^ isodecoders with the highest tRNA mature score promote SC-RT on UGA when overexpressed. Cys01 and Cys02 (**A**), Cys03, Cys04, and Cys05 (**B**), Cys06 – 09 (**C**) were overexpressed in HEK293T cells and SC-RT at UGA was measured using the p2luci system. Changes in the readthrough levels relative to the reporter with no tRNA (eV) were analysed by the Student’s t-test (mean+SD; n=2); statistically significant increase with P < 0.01 is marked with asterisk “*”.

Finally, we employed the pSGDluc system and observed that the UGA SC-RT values were smaller and varied to a lesser extent compared to those measured in p2luci. We observed statistically significant increase of SC-RT on UGA (P < 0.01) when Cys01, Cys03 and Cys06 were overexpressed (Fig. 2; for numbers see Supplemental Excel File 1). The highest impact of Cys01 is consistent with the p2luci results.

**Figure 2.**
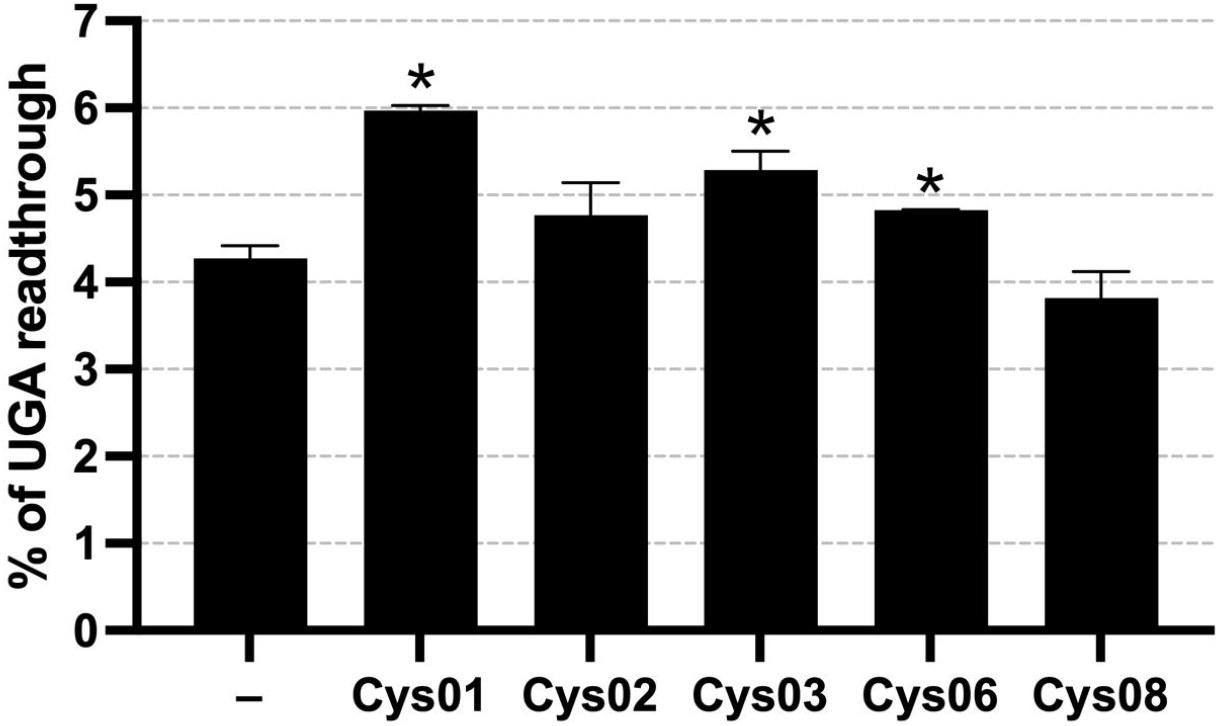
Cys01 displays the highest readthrough levels at UGA also with the pSGDluc system. The changes in the readthrough levels relative to the reporter with no tRNA (eV) were analysed by Student’s t-test (mean+SD; n=2); statistically significant increase with P < 0.01 is marked with asterisk “*”.

### Sequential and structural differences of the tRNA^Cys^ isodecoders

Cys01, with the second highest mature tRNA score of all the tRNA^Cys^ isodecoders (81.8; Supplementary Table 1), repeatedly showed the highest UGA SC-RT values in both reporter systems. It was also the second highest ranking isodecoder in ChIP-Seq data analysis (Supplementary Table 3) and the third highest ranking isodecoder in YAMAT-Seq (Supplementary Tables 4 - 5). The sequence and the secondary structure of this isodecoder appears to be canonical, showing no unusual residues or introns (Fig. 3A). Owing to its highest SC-RT potential, the Cys01 sequence and structure was taken as a reference for comparison with other isodecoders in order to identify nucleotides with the highest impact on SC-RT of tRNA^Cys^.

**Figure 3.**
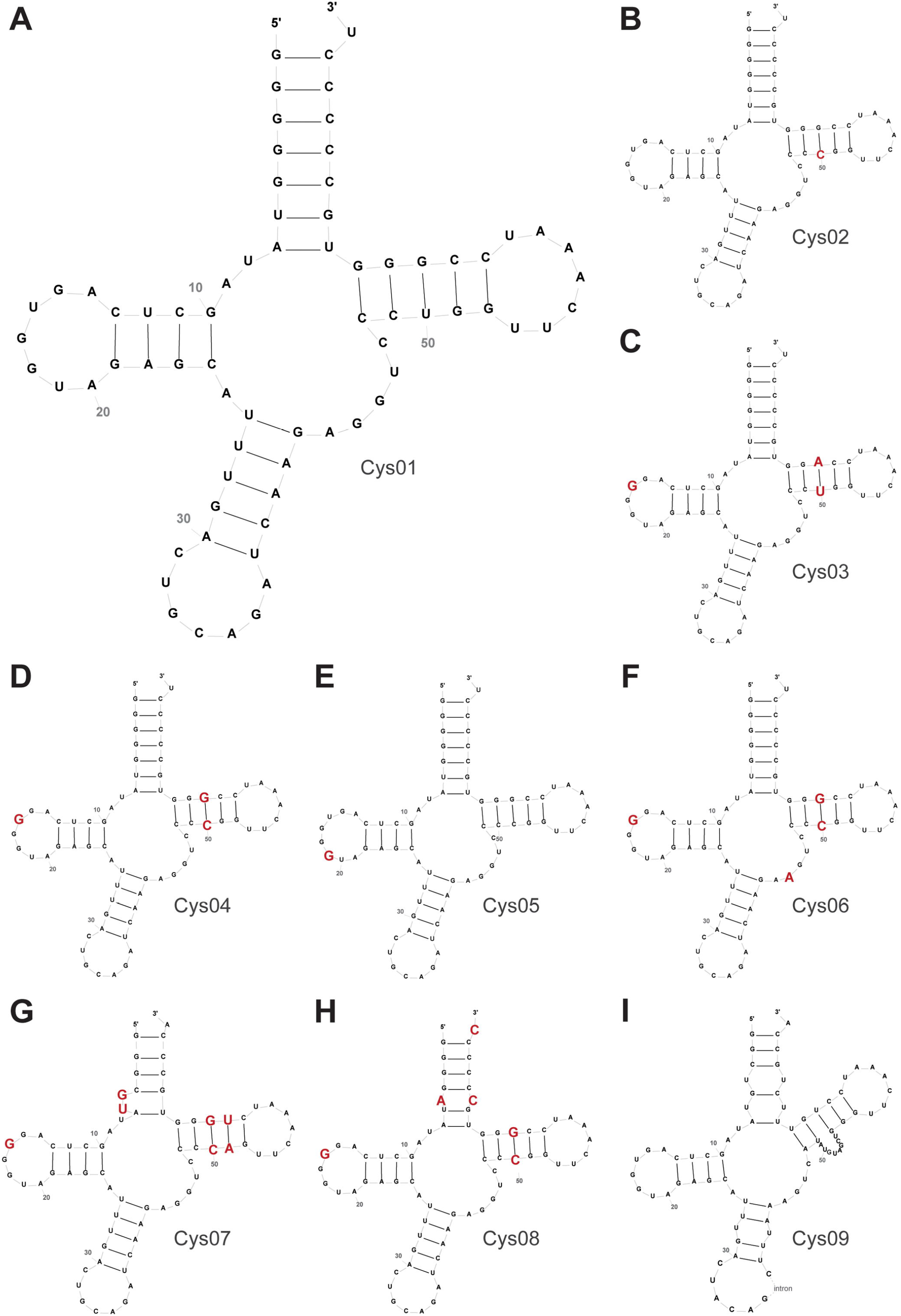
Comparison of primary sequences and secondary structures of tRNA^Cys^ isodecoders. Cys01 (**A**) was taken as a reference sequence /structure - all individual differences between Cys01 and other isodecoders (**B – I**) are highlighted in red. The secondary structure predictions were done with help of http://gtrnadb.ucsc.edu.

Cys02, with the highest mature tRNA score (81.9; Supplementary Table 1), displayed the second highest UGA SC-RT values with p2luci. Remarkably, it was also the highest ranking isodecoder in all three expression analyses (Supplementary Tables 2 - 5). Note that human genome contains four identical gene copies of Cys02; tRNA-Cys-GCA-2-1, tRNA-Cys-GCA-2-2, tRNA-Cys-GCA-2-3 and tRNA-Cys-GCA-2-4.

Compared to Cys01, there is only one difference in the Cys02 sequence; cytosine instead of uracil 49 in the 3^rd^ base-pair of the T-arm turning this non-Watson-Crick G:U base-pair of Cys01 into a canonical G:C pair in Cys02 (Fig. 3B). This change seems to have only a minor effect (and only in pSGDluc) on SC-RT values.

The third highest UGA SC-RT values were obtained with Cys03, which scored in all three expression analyses lower than the former two tRNAs (Supplementary Tables 2 - 5). Compared to Cys01, the 3^rd^ base-pair (G49 : U61) of the T-arm changed to A:U and uracil 16 in the D-loop was replaced with guanine (Fig. 3C). Interestingly, Cys04, showing modestly but still significantly increased UGA SC-RT values, and reaching quite low numbers in all the three expression analyses (Supplementary Tables 2 - 5), carries G:C as the 3^rd^ pair of the T-arm, like Cys02, however, contains U16G substitution in the D-loop, like Cys03 (Fig. 3D). Since Cys05, with no significant SC-RT activity, also carries G:C as the 3^rd^ pair of the T-arm but contains an extra G (19) in the D-loop (Fig. 3E), we conclude that the composition of the D-loop most probably determines the efficiency of SC-RT of tRNA^Cys^ isodecoders).

In accord, Cys06 with the U16G substitution in the D-loop, carrying also the G:C as the 3^rd^ pair of the T-arm and an extra G44A substitution in the center body of tRNA, fails to promote SC-RT (Fig. 3F). Interestingly, in ChIP-Seq analysis, Cys06 displayed relatively high expression levels (about +++; Table 2), in fact much higher than those of Cys04 or Cys05. Similarly, Cys07 and Cys08 with the U16G substitution in the D-loop, G:C as the 3^rd^ pair of the T-arm and additional changes elsewhere were not active in SC-RT (Fig. 3G and H), yet Cys08 was the second highest ranking isodecoder in tRNA-Seq data from Torres (2019) (Supplementary Table 2) and the top runner in the ChIP-Seq analyses (Supplementary Table 3). An exceptional case is Cys09 carrying a non-canonical variable loop with no activity either (Fig. 3I). Note that Cys09 also contains an intron between sequence positions 37 and 133 and is categorized as a pseudogene in GtRNAdb.

These results also suggest that while the mature tRNA score is a valuable predictor of SC-RT-promoting activity of tRNAs, it cannot used as a predictor of their stable expression and general functionality. On the other hand, because we are unable to reliably quantify the level of overexpression of our tRNAs for the technical reasons described previously (Beznoskova et al. 2021), we also cannot say with 100% certainty that tRNAs that do not promote SC-RT are stably expressed from our reporters and properly aminoacylated. In other words, we can be certain of those tRNAs that show SC-RT stimulation, but cannot completely rule out that those that are known to be expressed but failed to promote SC-RT in our assays are not well expressed in our systems. This caveat, however, does not detract from our finding that at least some tRNA^Cys^ isodecoders function as rti-tRNAs in humans together with tRNA^Trp^ and tRNA^Tyr^, like in yeast Beznoskova, 2016 #5981;Beznoskova, 2019 #6302}. Whether or not also the fourth “yeast” rti-tRNA (tRNA^Gln^) fulfills the rti-tRNA role also in humans is currently being investigated in our lab.

## DISCUSSION

The aim of this study was to identify a functional tRNA^Cys^ molecule(s) capable of promoting SC-RT in human cells. Since all tRNAs in the GtRNAdb are only computationally predicted and it is not known which genes encode functional tRNAs, we analyzed three expression datasets acquired by different methods from different cell lines (Rudolph et al. 2016; Shigematsu et al. 2017; Torres et al. 2019). We reasoned that despite method-specific limitations, such as secondary structures and post-transcriptional modifications of tRNAs that may impair tRNA-seq sequencing, or Pol III stalling that may lead to false positive ChIP-Seq results, at least some approximation of *in vivo* expression levels should be considered in addition to pure bioinformatics. Cross-comparison of the expression levels of tRNA-Cys isodecoder genes among these datasets confirmed that the expression of tRNA isodecoders varies widely and not all tRNA isodecoder genes are expressed in all cells and tissues to the same extent (Kutter et al. 2011; Torres et al. 2019). We consider this fact, plus technical hurdles that prevent us from reliably quantifying the rate of overexpression of tRNAs^Cys^ in HEK293 cells ((Beznoskova et al. 2021) – we perform only transient transfections, the efficiency of which varies and we have no means to determine the number of non-transfected cells that would need to be accounted for), as a shortcoming of our study, because the actual levels of tRNA^Cys^ isodecoders in HEK293 cells are unknown and can be only approximated. However, this does not prevent us from achieving our goal; it only limits us in that we cannot draw clear conclusions about tRNAs other than those that achieved positive results in our assays.

The fact that the UGA SC-RT levels measured in p2luci differ from the levels measured in pSGDluc might be attributed to the fact that two different stop codon contexts have been used; the TMV context in p2luci and the –UAG-C context in pSGDluc. In support, differing SC-RT levels were described before in an experiment comparing pSGDluc to the traditional dual-luciferase vector system pDluc, which is similar to p2luci, with a distinct stop codon context (Fixsen and Howard 2010).

In p2luci, two molecules, tRNA-Cys-GCA-4-1 (Cys01) and tRNA-Cys-GCA-2 (Cys02) showed almost ∼ 100 % and ∼ 50 % increase in SC-RT levels, respectively, when overexpressed in HEK293T. Compared to all tested isodecoders with lower to none SC-RT promoting activity, the latter tRNAs differ in the 49:61 pair of the T-arm but other than that share exactly the same sequence including the D-loop (Fig. 3). All others, with the exception of the pseudogene Cys09, displayed variability not only in the T-arm but also in the D-loop. Therefore, taking into account the shortcoming mentioned above, we propose that the sequence of the D-loop, an in particular U16, critically contribute to the SC-RT-promoting ability of tRNAs^Cys^ by a mechanism that remains to be explored.

With respect to a potential role of tRNA modifications, there is no much to report because no human cysteine tRNA modifications can be found in the MODOMICS database. However, some modifications for tRNA-Cys were suggested by novel high-throughput sequencing methods, namely m^1^A58 and m^1^G37 modifications for both tRNA-Cys-GCA-2-2 and tRNA-Cys-GCA-4-1 isodecoders, using demethylase tRNA sequencing (DM-tRNA-seq) in HEK293T cells (Clark et al. 2016). However, none of these bases shows any variability among all tRNAs^Cys^ and, therefore, these modifications should make no difference. Notably, in case of tRNAs^Cys^, the GCA anticodon and U73 represent the general identity determinants for aminoacylation (Pallanck et al. 1992). Since also U73 does not differ among tRNA^Cys^ isodecoders, we assume that charging should also not make any difference.

Interestingly, we noticed that overexpression of Cys04, Cys05, Cys07, and Cys09 in the p2luci system resulted in a partial translational shut down, which could be explained by generation of the 5’ tRNA-derived fragments. However, more experiments are needed to prove or disprove this idea.

Taken together, we conclude that, in agreement with our previous data with tRNA^Tyr^ and tRNA^Trp^ (Beznoskova et al. 2021), the “tRNA score” is a reliable predictor of a functionality of a given tRNA iso-decoder in SC-RT, but not necessarily of its stable expression. In any case, this study extends the aforementioned repertoire of human rti-tRNAs to include also rti-tRNA^Cys^.

## MATERIALS AND METHODS

### Plasmids

The lists and descriptions of plasmids, primers, and GeneArt™ Strings™ DNA Fragments used throughout this study (summarized in Supplementary Tables 7 – 9) can be found in the Supplementary Information. The tRNA sequences were obtained by tRNAscan-SE Analysis of *Homo sapiens* (hg19 - NCBI Build 37.1 Feb 2009) (Lowe and Eddy 1997).

### Cell lines manipulations

Hek293T cells (human embryonic kidney cells obtained from Petr Svoboda, IMG of Czech Academy of Sciences) were cultured in DMEM (Sigma, cat # D6429) supplemented with 10% fetal bovine serum FBS (Sigma, cat # F7524). The H1299 cells were cultured in RPMI plus GlutaMAX (Gibco by Life Technologies) supplemented with 10% fetal bovine serum (FBS, Gibco by Life Technologies) and 100U/ml of penicillin/streptomycin. Cells were kept in a humidified atmosphere containing 5.5% CO_2_ at 37°C.

### Stop Codon Readthrough Assays

For one reaction, ∼40,000 cells were plated into a single well of a 24-well plate and 24 hours later transfected with 200ng of DNA (for UAG, UGA and sense control reporters) or 500ng of DNA (for UAA) using Turbofect (Invitrogen). No cell toxicity was observed for these DNA doses. Please note that when using a 6-well plate, 5-times more of each component must be taken into each reaction. A properly mixed reaction (100µl of DMEM (Sigma) with DNA and Turbofect (1µl per 500ng of DNA)) was incubated at the room temperature for 20 minutes and then applied onto the cells. SC-RT was assayed 24 hours after the transfection as follows. The media mixture was removed from each well and the cells were lysed with a room tempered 1xGlo buffer (Promega) for 5 min at 25°C while shaking at 550 rpm. Equal amounts of the lysate were transferred into two Microtiter plates (one for Renilla and the second for Bright Glo (Promega)). Substrates for both luciferases were added in a one-to-one ratio and the reactions were incubated for 5 minutes (BrightGlo) or 15 minutes (Renilla) at 25°C while shaking at 225 rpm. The luminiscence was measured by Tecan Infinite 200Pro. The SC-RT efficiency was estimated by calculating the ratio of Firefly luciferase to Renilla luciferase activity obtained with the test constructs and normalizing it to the ratio obtained with an in-frame control construct. For each construct with two or three reactions at the time; at least five independent transfection experiments were performed.

## ACKNOWLEDGMENTS

We are thankful to all lab members for fruitful discussions. This work was supported by the Czech Science Foundation grants 20-00579S and 23-08669L and the Praemium Academiae grant provided by the Czech Academy of Sciences (all to L.S.V.).

## SUPPLEMENTARY DATA

Supplementary Data are available at The RNA journal online.

## Notes

### Competing Interest Statement

The authors have declared no competing interest.

